# Increasing Signal Intensity of Fluorescent Oligo-Labeled Antibodies to Enable Combination Multiplexing

**DOI:** 10.1101/2023.07.06.547965

**Authors:** Madeline E. McCarthy, Xiaoming Lu, Oluwaferanmi Ogunleye, Danielle R. Latham, Megan Abravanel, Daniel Pritko, Jonah R. Huggins, Charlotte V. Haskell, Nishi D. Patel, Zachariah A. Pittman, Hugo Sanabria, Marc R. Birtwistle

## Abstract

Full-spectrum flow cytometry has increased antibody-based multiplexing, yet further increases remain potentially impactful. We recently proposed how fluorescence Multiplexing using Spectral Imaging and Combinatorics (MuSIC) could do so using tandem dyes and an oligo-based antibody labeling method. In this work, we found that such labeled antibodies had significantly lower signal intensity than conventionally-labeled antibodies in human cell experiments. To improve signal intensity, we tested moving the fluorophores from the original external (ext.) 5’ or 3’ end-labeled orientation to internal (int.) fluorophore modifications. Cell-free spectrophotometer measurements showed a ∼6-fold signal intensity increase of the new int. configuration compared to the previous ext. configuration. Time-resolved fluorescence and fluorescence correlation spectroscopy showed that ∼3-fold brightness difference is due to static quenching most likely by the oligo or solution in the ext. configuration. Spectral flow cytometry experiments using peripheral blood mononuclear cells show int. MuSIC probe-labeled antibodies (i) retained increased signal intensity while having no significant difference in the estimated % of CD8+ lymphocytes and (ii) labeled with Atto488, Atto647, and Atto488/647 combinations can be demultiplexed in triple-stained samples. The antibody labeling approach is general and can be broadly applied to many biological and diagnostic applications where spectral detection is available.

## Introduction

Fluorescent antibodies are an integral tool for biological and diagnostic applications^1^. One application for fluorescent antibodies is flow cytometry^2^. The use of fluorescent antibodies with conventional flow cytometers is restricted to typically 3-4 markers, but up to ∼10-15 markers have been reported^2–4^. The restriction is largely due to spectral overlap between fluorophores, limiting the number of analytes that can be reliably detected. Regardless, flow cytometry remains a useful platform as it is a cost-effective, high-throughput, and non-destructive method for single-cell analysis^5,6^. Recent advances have led to full-spectrum flow cytometry (FSFC), which captures the entire fluorophore emission spectra, creating a unique spectral fingerprint for each fluorophore^7,8^. This allows fluorophores with similar peak emissions to be used in the same panel, so long as they have distinctive spectral signatures. FSFC has enabled the simultaneous detection of up to 40 markers^9^, but further multiplexing capabilities are stunted by the number of commercially available dyes compatible within a single panel. Moreover, FSFC is still far from the multiplexing capabilities of methods such as single-cell RNA sequencing, which has the ability to identify 100s-10,000s of markers^10,11^.

The 40-plex FSFC panel largely relies on single-dye fluorescent antibodies, with relatively few tandem-dye fluorescent antibodies^9^. We recently developed Multiplexing using Spectral Imaging and Combinatorics (MuSIC), which uses combinations of currently available fluorophores to create spectrally unique MuSIC probes^12^. MuSIC probe-labeled antibodies may expand the multiplexing capability for FSFC by providing new tandem probes. We proposed an oligo-based method for covalently labeling antibodies with MuSIC probes **(Fig 1A-B)** and validated this method using spin column purification, absorbance measurements, and Protein A beads / spectral flow cytometry^13^, based on a previous protocol^14^.

**Figure 1.**
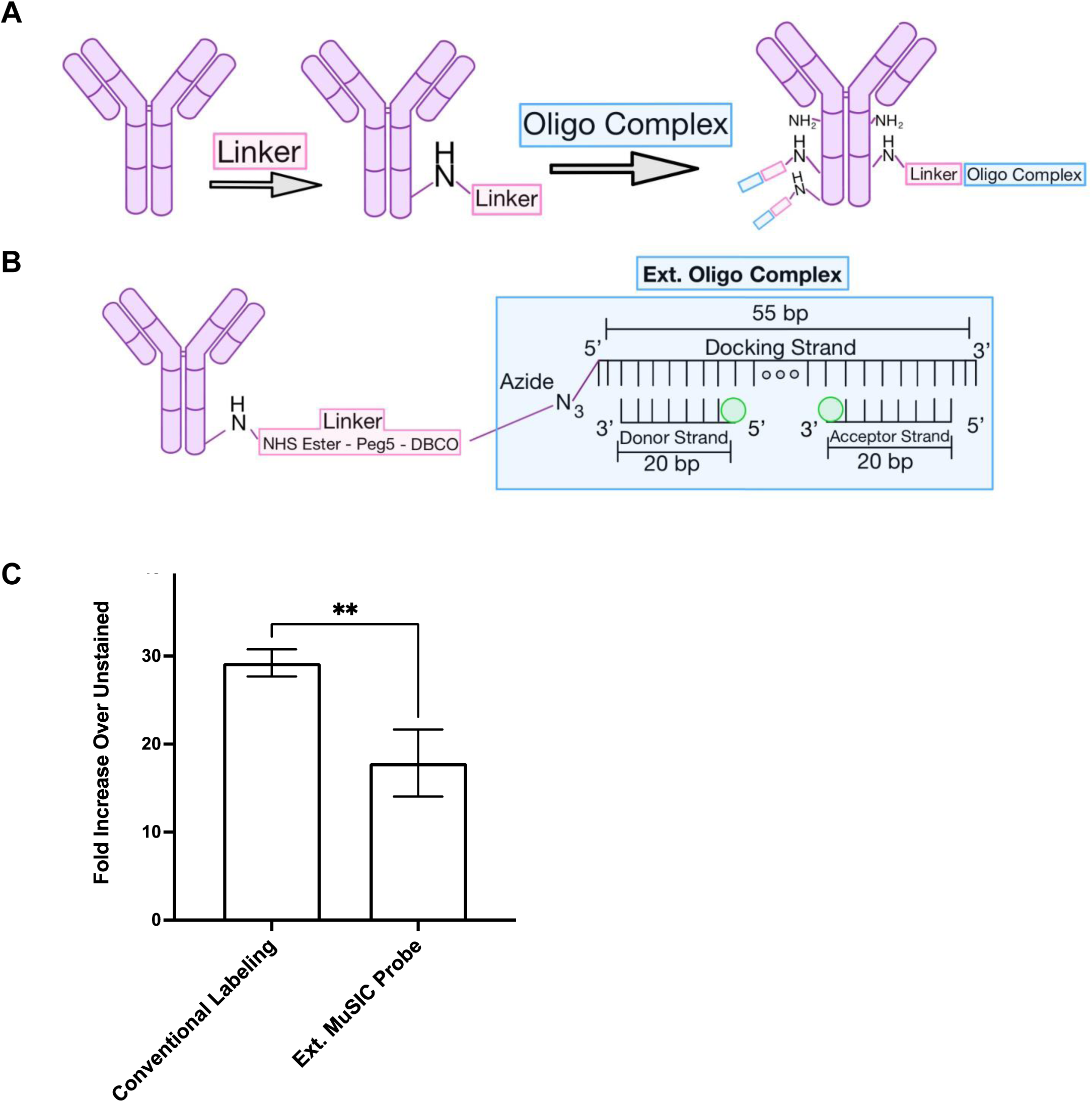
Oligo-based ext. MuSIC probe labeling of antibodies. (A-B) Graphic depicting MuSIC probe labeling. The NHS ester of the linker reacts with free amines on the antibody. Fluorophore-end-labeled (external-ext.) donor and acceptor strands are annealed onto the docking strand to form the oligo complex. The azide on the docking strand, in the oligo complex, is reacted with the free DBCO group on the linker to covalently bind the oligo complex to the antibody. There are multiple free amines on each antibody, allowing for the linker to attach at multiple sites, increasing the degree of labeling. (C) Comparison of fluorescence intensity of PBMCs stained with CF488A conventional labeling kit vs Atto488 ext. MuSIC probes. Error bars are standard error from triplicate measurements.

There are a variety of methods for conjugating oligos to antibodies. Some involve non-covalent complexation between an antibody and oligonucleotide through a unique linker, such as protein A/G–Fc or streptavidin–biotin^15^. These sets of reactions are characterized by, for example, the conjugation of maleimide-activated streptavidin to thiol-reduced oligonucleotide to create a streptavidin-oligonucleotide substrate which further undergoes a non-covalent coupling with a biotinylated antibody^16^. Yet others involve alternative covalent linking chemistries, such as the coupling between reduced antibody and maleimide-activated oligonucleotide^17^. While some of these are not incompatible with our approach, our developed method was shown to allow placement of fluorophore pairs with some control over the shape of tandem probe spectra, hence our continued focus there. However, our method is yet to be tested on human cells.

In the current study, we first applied our previous method to staining human peripheral blood mononuclear cells (PBMC). In doing so, we found significantly lower staining intensity compared to a conventional antibody labeling kit (Biotium Mix-n-Stain), which would often cause analysis problems due to low signal-to-noise. Consequently, we hypothesized that a different oligo-fluorophore arrangement of the MuSIC probes, using internal fluorophore modifications rather than external fluorophore modifications, could increase the fluorescent signal intensity of MuSIC-probe labeled antibodies. Results showed that the new method with internal fluorophore modifications produced ∼6-fold increase in fluorescent signal compared to the previous method. Biophysical characterization showed that ∼3-fold of this difference is due to fluorescence static quenching, most likely induced by the oligo or solution environment, rather than by other fluorophores. We then compared the internally modified MuSIC-probe labeled antibodies to conventionally labeled antibodies by staining PBMCs. Results revealed that the new internal labeling method retained increasedsignal intensity while exhibiting no significant difference in the estimated percentage of CD8+ lymphocytes. Lastly, we demonstrate that three different antibodies labeled with either Atto488, Atto647, or an Atto488/647 combination can be effectively unmixed in triple-stained PBMCs analyzed by FSFC. This enhanced fluorescent signal and unmixing capability suggests the potential of MuSIC-probe labeled antibodies to complement to the existing capabilities of FSFC, by providing new spectrally unique fluorescent antibodies with comparable intensity. Such antibodies are not restricted to FSFC but could be useful for other biomedical applications such as tissue heterogeneity studies with immunofluorescence imaging when spectral detection is available.

## Results

We previously developed a method for labeling antibodies with combinations of fluorophores (i.e. MuSIC probes)^13^. In short, an oligo complex containing fluorescent molecules is conjugated to the antibody via a DBCO-Peg5-NHS ester (referred to as the linker) (**Fig 1A**). Here the oligo complex is composed of a 20 bp oligo with a 5’ end fluorophore modification (referred to as the 5’ donor strand) and a 20 bp oligo with a 3’ end fluorophore modification (referred to as the 3’ acceptor strand) that are co-hybridized to a 55bp oligo with a 5’ azide modification (referred to as the docking strand) to form the externally labeled (ext.) oligo complex (**Fig 1B**). This is called ext. because it refers to the method of fluorophore labeling on the donor and acceptor strands, which is external end labeling (to be contrasted later). These figures are not meant to convey distances or 3D versus 2D orientations, but rather to serve as illustrations.

We previously demonstrated covalent labeling of antibodies with MuSIC probes using this method and validated the labeling protocol with spin-column purification, absorbance measurements, and FSFC measurements with protein A beads bound to (i) Cy3, (ii) Tex615, and (iii) Cy3/Tex615 ext. oligo labeled antibodies^13^. Since this method had only been validated using beads, we asked whether this method would work when staining peripheral blood mononuclear cells (PBMCs). We created an ext. oligo complex using an Atto488 5’ donor strand and an Atto488 3’ acceptor strand as the MuSIC probe and conjugated it to anti-CD8 antibodies. For comparison, we used a commercially available Biotium Mix-n-Stain kit to conventionally label CD8 antibodies with CF488A dye, which is reported to have comparable fluorescent properties (excitation peak, emission peak, and brightness) to Atto488^18^. PBMCs were stained with each antibody batch and analyzed by FSFC. Results showed that the median signal intensity of cells stained with the ext. labeled MuSIC probe was ∼2-fold (p-value=0.01) lower compared to cells stained with conventionally labeled antibodies (**Fig 1C**). The lower intensity interfered with data analysis, in terms of assessing positively stained cells, therefore needed to be addressed to move forward with the approach.

We then asked how we can increase the signal intensity of MuSIC probe-labeled antibodies. We reasoned that the lower fluorescence signal was not due to the degree of labeling because it was previously calculated to be within the standard range^13,19^. Some degree of difference in signal intensity may be due to differences in dye properties between CF488A and Atto488, although as mentioned above, the dyes sharesimilar characteristics. To investigate whether the docking strand and/or hybridization played a role, we examined the fluorescence emission intensity of Atto488 5’ donor strands and Atto488 3’ acceptor strands alone in solution and when co-hybridized to the docking strand (**Fig 2A**). We observed that the hybridization of the 5’ donor and 3’ acceptor strands to the docking strand results in a significant decrease in fluorescent signal compared to the strands on their own. While one contribution may be self-quenching, using only the donor or acceptor showed similar trends, strongly suggesting the involvement of other mechanisms.

**Figure 2.**
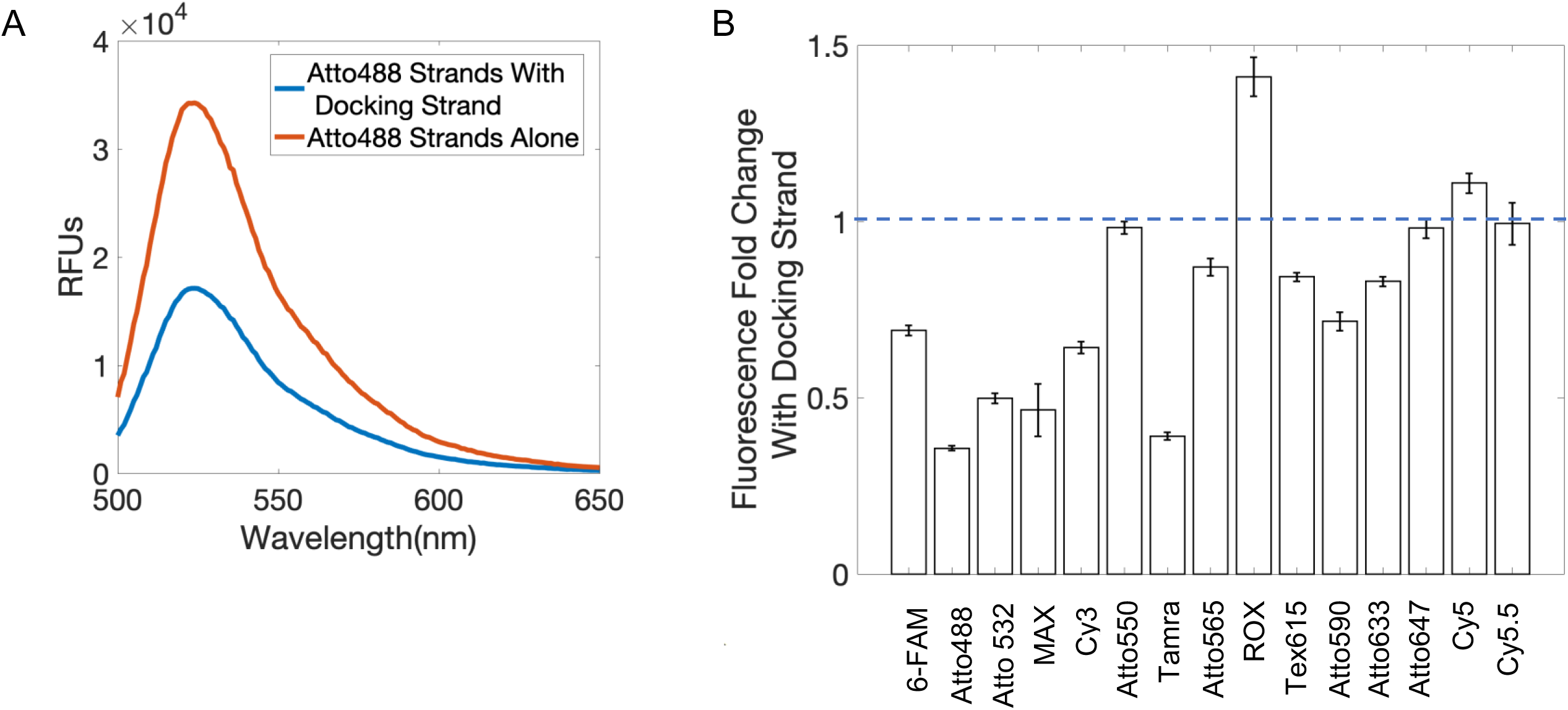
Fluorescence signal change from Docking Strand. (A) Comparison of fluorescence emission spectra, excited at 470nm, of the Atto488 5’ Donor and 3’ Acceptor strands hybridized to the Docking Strand and when alone in solution with and without the Docking Strand. Data are representative from triplicates. (B) Change in fluorescence intensity of 15 fluorescent oligos when hybridized to the Docking Strand. Error bars are standard error from triplicates.

We further wondered whether this docking strand-induced fluorescence decrease was a fluorophore-specific phenomenon or if it occurred for other fluorophores. Therefore, we compared the emission intensity with and without docking strand for 15 different fluorophore-conjugated 5’ donor strands and 3’ acceptor strands (**Fig 2B**). Signal decreased with docking strand for all but five of the fluorophore-conjugated strands that were tested. Although proximity-based quenching is a potential explanation for these observations, these experiments explored cases where only donor or acceptor strand were bound. Thus, even when there is no chance for quenching between fluorophores, similar decreases were observed. This suggests other mechanisms are driving the fluorescence decrease.

Previous studies showed that there can be a significant change in fluorescence when oligo-strands containing an end-fluorophore modification are hybridized to strands containing an overhang^20^, such as in the ext. oligo complex. Other studies have suggested fluorescence quenching can be universally mediated by water and other solvents^21^. These findings led us to hypothesize that if the fluorophores within the oligo complex had a different orientation, it could give an increased fluorescent signal. To test this, we adjusted the configuration of the ext. oligo complex to contain oligos with internally (int.) conjugated fluorophores, guided by a recent study creating a FRET ruler^22^. The int. nomenclature denotes the method of oligo labeling, which occurs internally within the oligo, to contrast with ext. end labeling. This new oligo complex consists of the 50 bp int. acceptor strand and a 15 bp azide strand which both co-hybridize to the 65 bp int. donor strand (**Fig 3A**). The purpose of a separate azide strand is to reduce the cost of oligo production, due to the increased difficulty of synthesizing an oligo with two modifications. The new donor and acceptor strands both have an internal fluorophore modification (int donor and int acceptor), rather than 5’ and 3’ end fluorophore modification, respectively. We created int. and ext. oligo complexes (both using Atto488 conjugated strands) and measured their fluorescent emission spectra. We observed a ∼6-fold fluorescent signal increase of the int. oligo complex compared to the ext. oligo complex in solution (**Fig 3B**). Note that the distance between fluorophores is expected to be similar in both cases, so the removal of quenching between fluorophores is expected to not fully explain the increased fluorescence intensity.

**Figure 3.**
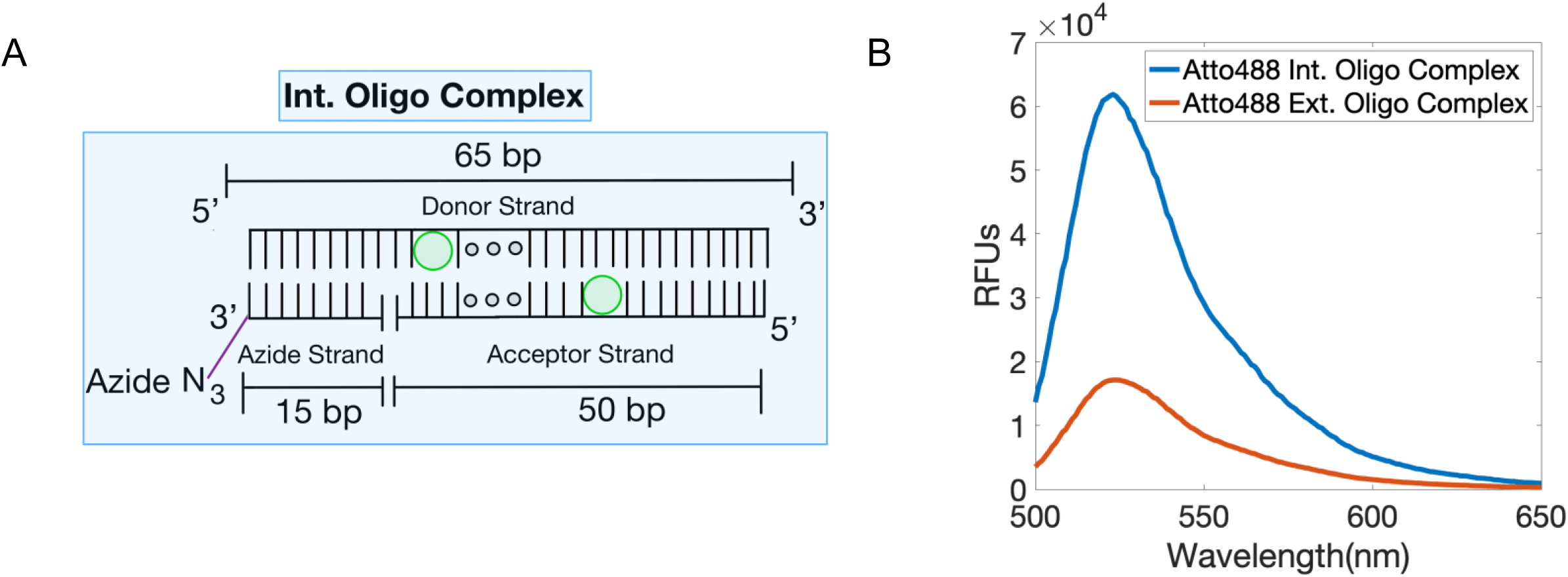
The int. labeling method increases intensity relative to the ext. labeling method. (A) Int. oligo complex containing the Int. Donor and Acceptor strands and the Azide strand. (B) Comparison of relative fluorescence intensity of the Atto488 probes using the int. and ext. oligo complexes (470 nm excitation). Data are representative from triplicates.

Before functional testing, we wanted to better understand the nature of the fluorescent signal differences between the int. oligo complex and ext. oligo complex. We used Time Resolve Fluorescence Spectroscopy (TRFs) to investigate the quenching mechanism and rotational correlation time, as well as Fluorescence Correlation Spectroscopy (FCS) to derive the number of bright molecules, the translational diffusion, and the triple state kinetics (**Fig 4**). We found that the ext. oligo complex undergoes more dynamic quenching than the int. oligo complex, as shown by a shorter fluorescence decay of the ext. oligo, which can also be quantified using the species average lifetime (**Fig 4A**, **Table 3**). Dynamic quenching alters the average fluorescent decay lifetime by acting on the entire excited-state fluorophore population through rate dependent processes such as diffusion^23^. The ext. oligo complex also spends more time in the dark triplet state than the int. oligo complex, as shown by the differences in the correlation curves (**Fig 4B**, **Table 3**). Further, since FCS only monitors bright molecules, a single (bright) int. oligo complex is only 1.6 times brighter than an ext. oligo complex (**Table 3**). The observed ∼6-fold difference in intensity (**Fig. 3B)** but < 2-fold difference in the molecular brightness (**Table 3**) of the int. oligo complex relative to the ext. oligo complex can be explained through the formation of a static ground-state complex which renders fluorophores non-observable, commonly referred to as static quenching^23^. Hence, we conclude that ∼3-fold difference in intensity is due to static quenching in the ext. oligo complex. However, this does not reveal whether quenching between fluorophores or between fluorophores and the environment (oligo or solution) are responsible; based on the above observations we expect the latter to be more predominant. As expected, both oligo complexes show similar diffusion and rotational correlation times **(Fig 4B and C)**.

**Figure 4.**
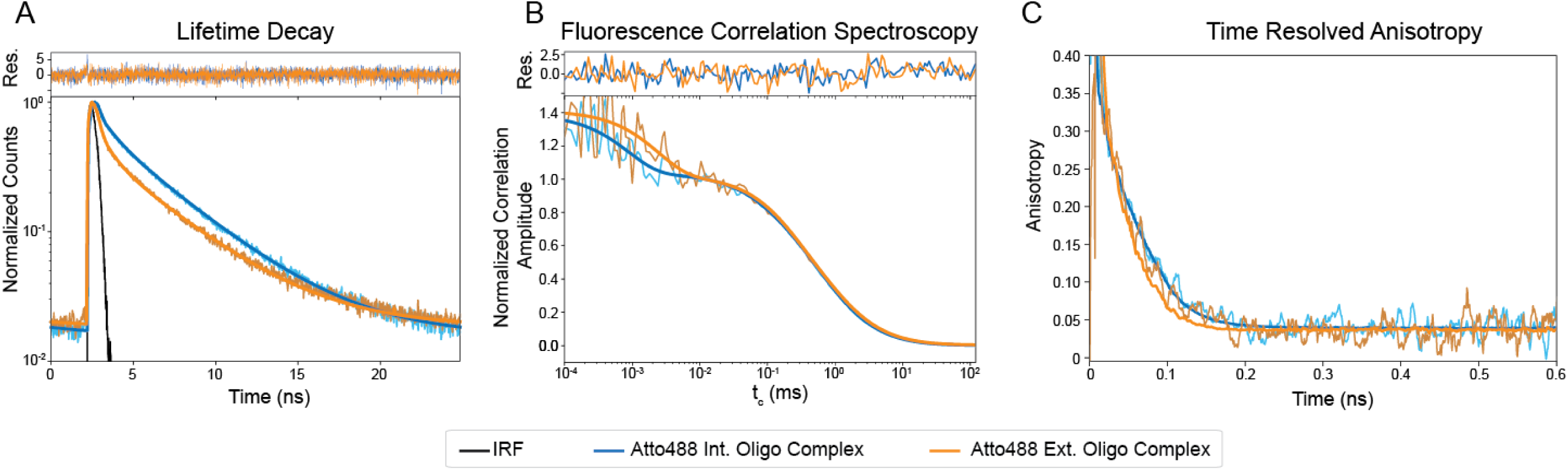
Differentiating static and dynamic quenching by Time-Resolved Fluorescence and Fluorescence Correlation Spectroscopy. Atto488 int. oligo complex fitting is shown in blue, with the raw data in light blue and Atto488 ext. oligo complex fitting is shown in orange, with the raw data in light orange. (A) Normalized fluorescence decays between Atto488 int. oligo complex and Atto488 ext. oligo complex. The difference in fluorescence lifetimes is visible by the difference in the slope of the decays. Residuals for the fitting model are shown on top. (B) Normalized fluorescence correlation between Atto488 int. oligo complex and Atto488 ext. oligo complex. The difference in dark triplet states is visible in the offset between the start of the curves to correlation times (t_c_) ∼10^-3^ ms. The curve overlaps between t_c_ being 10^-1^ and 10, indicating similar diffusion coefficients between samples. Residuals for the fitting model are shown on top. (C) Anisotropy rotational correlation between Atto488 int. oligo complex and Atto488 ext. oligo complex. The slight offset in rotational times could be due to differences in flexibility of the int. and ext. oligo complexes.

With this increase in signal intensity, we then asked how new int. MuSIC probe-labeled antibodies would compare to conventionally labeled antibodies when staining PBMCs for estimation of specific cell type abundances. Similar to above, int. oligo complexes with Atto488 were conjugated to CD8 antibodies to create int. MuSIC probe-labeled antibodies and CF488A was conjugated to CD8 antibodies using a Mix-n-stain kit to create the conventionally labeled antibodies. As above, while we cannot make definitive claims as to relative signal intensity between the two due to differences in labeling chemistry and fluorophore type, the fluorophores are highly similar, and the intent is not to directly claim these int. oligo labeled antibodies are brighter. PBMCs were stained with each antibody batch and analyzed by FSFC. The signal-to-noise ratio (SNR) of cells stained with the int. labeled MuSIC probe was ∼10 fold higher compared to cells stained with conventionally labeled antibodies (**Fig. 5A**). We found no significant difference between the int. MuSIC probe-labeled antibodies and conventionally labeled antibodies for the % of CD8+ lymphocytes detected (**Fig 5B**). These results demonstrate that we were able to improve the design of MuSIC-probe labeled antibodies to increase the signal-to-noise ratio, with staining behavior comparable to conventionally labeled antibodies.

**Figure 5.**
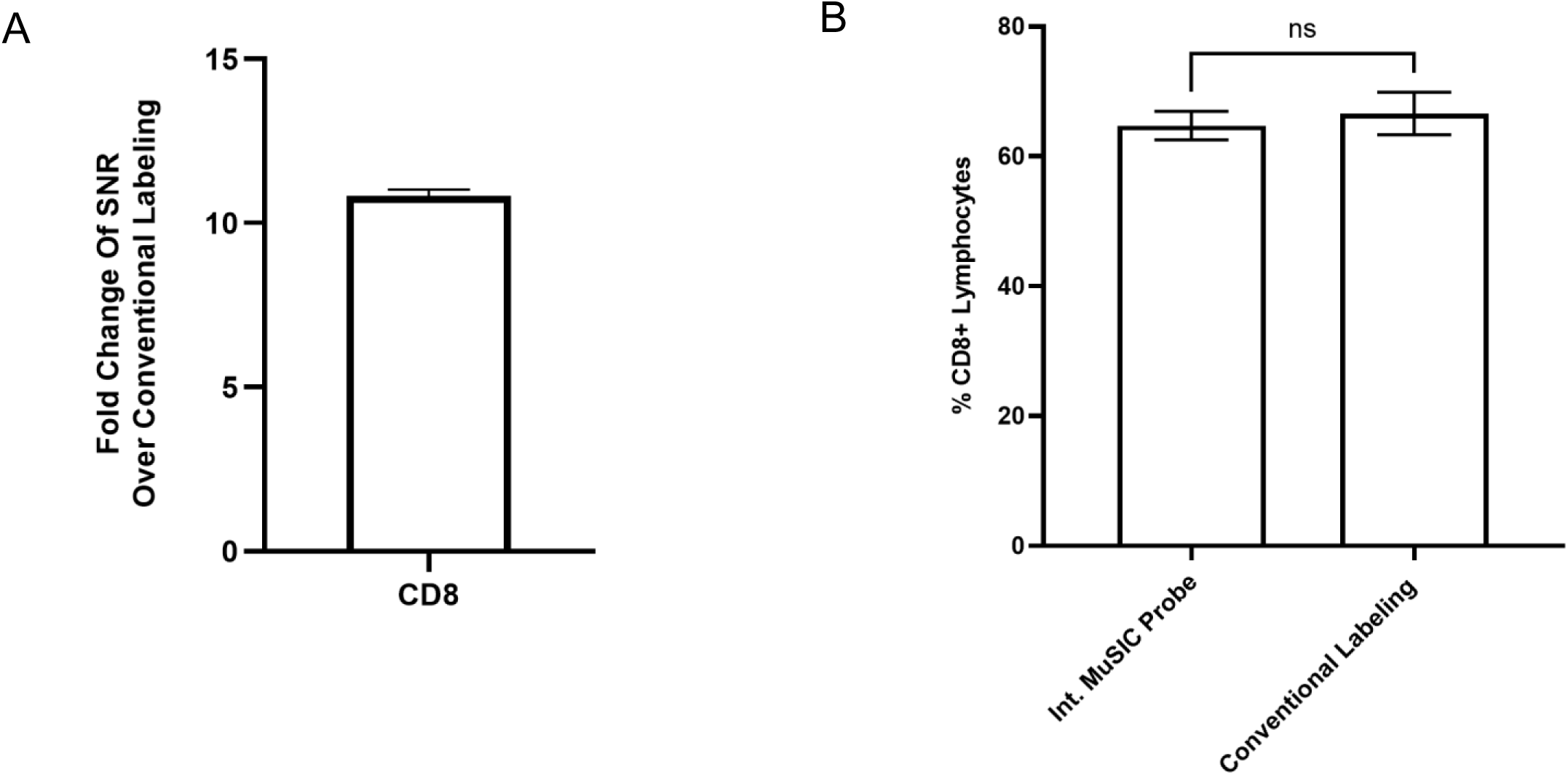
Comparing int. oligo complex and conventionally labeled antibodies in cell-based experiments. (A) Signal increase of PBMCs stained with Atto488 int.-labeled CD8 antibodies versus CF488A conventionally-labeled CD8 antibodies. (B) Percentage of CD8+ lymphocytes in PBMC for int.-labeled CD8 antibodies compared to CF488A conventional-labeled CD8 antibodies. Measurements are in triplicate and error bars are standard error.

Finally, we wondered whether int. labeled antibodies could be multiplexed using the MuSIC combination approach. We labeled three different antibodies (anti-CD8, -CD3 and -CD4, respectively) with three different oligo complexes, Atto488, Atto647, or an Atto488/647 combination (**Fig. 6A-emission spectra in Fig. S1**). We first analyzed data where PBMCs were singly-stained with these antibodies and assayed with FSFC (**Fig. 6B-C**). To our surprise, despite quite unique emission spectra between the different probes (**Fig. S1**), the unmixing results using internal manufacturer software revealed difficulty separating the Atto488/647 probe even in these single-stained experiments (**Fig. S2—**substantial Atto647 reported in Atto488/647-stained cells). Because this software was a black box solution, we decided to export the raw numerical data and apply our own unmixing analysis (see Supplementary Code). These analyses showed that for each cell, the observed fluorescence intensity strongly correlated with the unmixed signal output (**Fig. 6B**), and that the unmixed signal in singly-stained cells was predominantly assignable to the only antibody that was used for staining (diagonal versus off-diagonal in **Fig. 6C**—similar trends were observed in triplicates). Thus, it was sensible to use this unmixing strategy to analyze triple-stained PBMC. We did so and compared the % positive cells for each of the antibodies used for single-stained versus triple-stained analyses, which showed generally good correspondence (there was a discrepancy in CD4-stained samples that was statistically significant but not alarmingly different). We conclude that the int. oligo-labeled antibodies enable multiplexed analysis of cells using full spectrum flow cytometry with combination tandem dyes designed via MuSIC principles.

**Figure 6.**
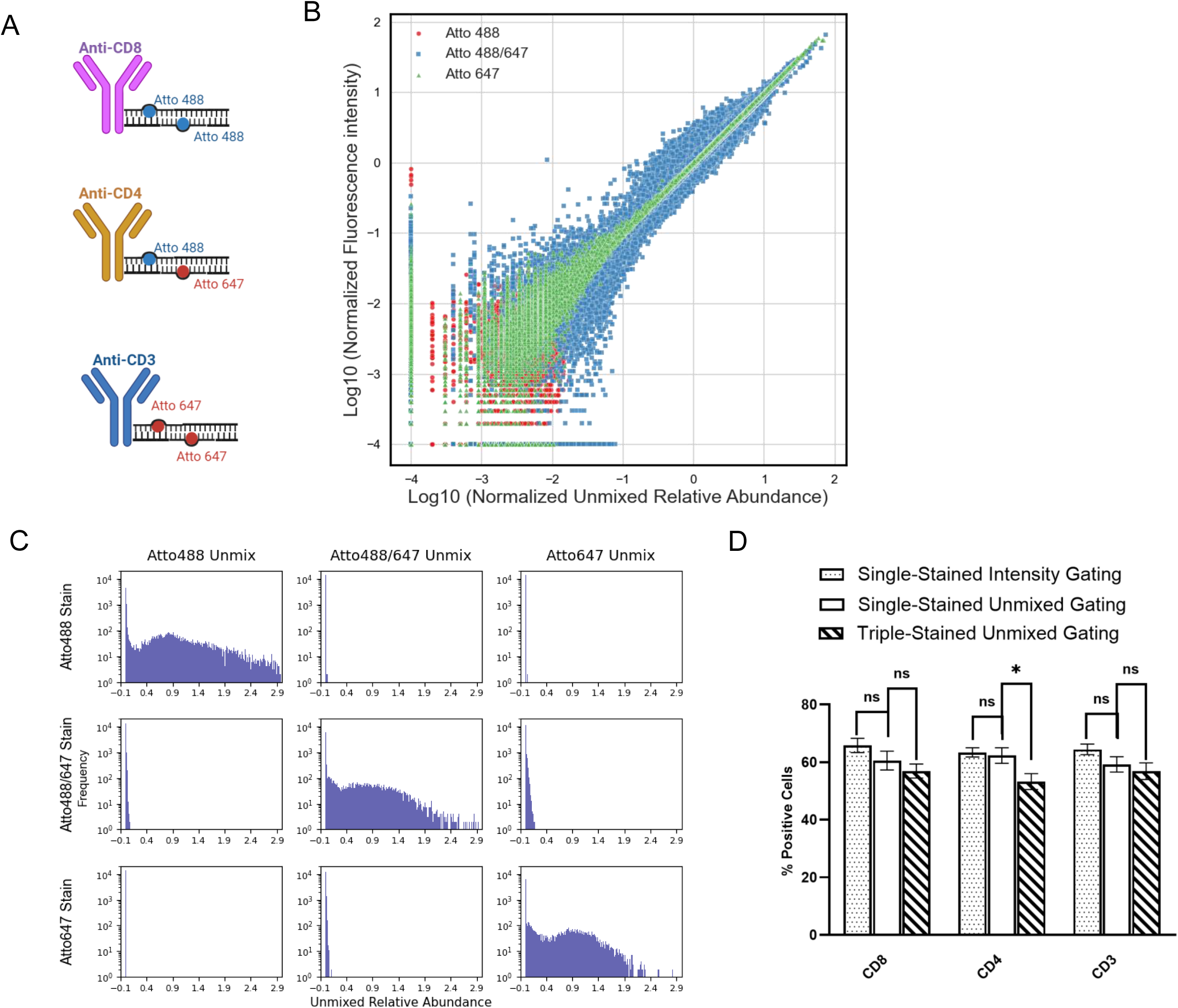
Unmixing Experiments for Int. Oligo-Labeled Antibodies. (A) Three different antibodies were labeled with three different oligo complexes, Atto488, Atto647, and the Atto488/647 combination. (B) The median fluorescence intensity (MFI) of positive cells in each group was used to generate the spectrum and identify the peak channel. Each cell’s fluorescence value at this peak channel was normalized by dividing it by the group’s MFI at the same channel. Similarly, the unmixed value of each cell was normalized by dividing it by its corresponding median value to generate the normalized unmixed relative abundance. The normalized fluorescence intensity in each cell is plotted versus its normalized unmixed relative abundance from PBMC singly stained with each antibody, demonstrating a good correlation between intensity and unmixed values. (C) Data exported from singly-stained PBMC were subjected to unmixing and histograms observed, showing good specificity. (D) PBMC were gated for positivity for the three different antibodies from histograms from either standard intensity-based metrics or unmixed quantification, in either singly-stained or triple-stained cell samples. There is overall good agreement between all methods demonstrating confidence in the approach. In all panels, measurements are based on triplicates and error bars are standard error. *: p<0.05 (t-test)

## Discussion

Here, we established a method to conjugate two fluorophores to an antibody and stain human cells with an increased signal intensity compared to our previous method. This method builds on our previously established labeling protocol but introduces key modifications to the oligo-fluorophore arrangement of the MuSIC probe. By re-arranging the oligo complex to eliminate the use of 3’ or 5’ end fluorophore modifications, we observe a significant increase in fluorescent signal for Atto488. Given the observed prevalence of docking strand-induced signal decrease, we expect this result may often apply to other fluorophores. We used Time Resolve Fluorescence and Fluorescence Correlation Spectroscopy to compare the old ext. oligo complex to the new int. oligo complex, finding that increased dynamic quenching and time in the dark triplet state explains the decreased fluorescence intensity of the ext. oligo complex. Multiple observations suggest the intensity change is not explained by simple fluorophore to fluorophore quenching. Using the new design, we stained human PBMCs and compared the signal intensity to that of conventionally labeled fluorescent antibodies, and observed a statistically significant increase in the resulting fluorescent signal without creating any significant differences in the % of CD8+ lymphocytes. We then demonstrated unmixing of three labeled antibodies with the approach using two fluorophores (Atto488 and Atto647) and their combination (Atto488/647), consistent with the MuSIC approach.

To maximize the potential of this new increased intensity probe design, the next step will be to select different combinations of fluorophores to assemble a palette of spectrally unique antibody-conjugated MuSIC probes. Here, we studied a simple Atto488 and Atto647 pair, because significant prior work with this pair in Förster Resonance Energy Transfer (FRET) rulers in oligos had already been done^22^. Approaches to expand can include stimulation studies for compatibility using a workflow similar to that described in our previous work^24^, and then testing the highest-ranked fluorophore combinations experimentally. For these simulations, the emission spectra of each potential MuSIC probe is generated, and lists of MuSIC probes that are likely to be deconvolvable in a mixture are generated and ranked. This ranking provides prioritization for testing experimentally by measuring the emission spectra of mixtures of MuSIC probes and unmixing them to determine which MuSIC probes can be accurately demultiplexed.

One major application of using MuSIC probe-labeled antibodies with FSFC can be cell-type profiling, which is the process by which a complex mixture of cell types, for example, from blood or tumors, are classified into the fractional composition of its components (e.g., neutrophils, natural killer cells, various types of T and B cells, etc.), based on classification of expression patterns (e.g., CD3 expressed or not)^25^. While there are 40 FSFC dyes available, very few of them are tandem dyes that can be used as uniquely identifiable markers, which limits the number of individual analytes that can be classified simultaneously. However, MuSIC probe-labeled antibodies could be used to expand the number of markers that can be detected by creating new combination fluorophore probes from the current dyes, to enhance current cell-type profiling efforts. FSFC has been previously paired with cell-type profiling to investigate the correlation between CD38 expression in macrophages and the predicted immune response to immune-checkpoint blockade therapy for hepatocellular carcinoma^26^. With a larger palette of compatible fluorescent tags, cell-type profiling efforts could expand further to look at an increased number of cell-type markers, for a more comprehensive view of a patient’s immune response to various treatments, or to complement other single cell profiling efforts^27–29^.

Additionally, MuSIC probe-labeled antibodies can be applied to a broad range of biological and diagnostic applications that involve the detection of protein expression. One of these applications can be for tissue imaging, such as recent highly multiplexed efforts^30–35^. If MuSIC probe-labeled antibodies can be combined with spectral imaging, this could allow for highly multiplexed, quantitative tissue imaging. One potential application is cancer, where increasing multiplexing capabilities could improve diagnostic potential by allowing for more tumor markers to be analyzed, thus leading to an increased mapping of tumor heterogeneity^36^. This could impact tumor detection, diagnosis, and treatment.

Although here we focused on increasing the fluorescent signal of oligo-based probes, by titrating the fluorescent oligos, we can decrease the fluorescent signal to a desired level in a controllable manner. Tunable fluorescence intensity is useful; for example, in static light scattering experiments^37^, where the sensitive photodiode detectors are easily saturated. Here, they labeled BSA at varying concentrations of fluorescent oligos between 0.03 - 0.10 µM that fluoresced below the saturation limit of the detectors while still achieving desired intensity. Conventional labeling kits would have been too intense, and as most are single reaction use, they can be difficult to control compared to the reported oligo-based probes which offer the unique advantage of reduced, tailorable intensities. In their case, the intensity tunability of the probes enabled a more flexible experimental design capable of separating simultaneous fluorescence and light scattering signals. The tunability of oligo-based fluorescent probe intensity could also be beneficial for cell staining, where some epitopes may have such a high abundance that a reduced fluorescent signal is necessary.

In addition to tunability for probe fluorescence intensity, the new int. oligo arrangement of these probes offers modulation of FRET between fluorophore combinations on the donor and acceptor strands. By adjusting the distance (bp) between the two fluorophores, one can increase or decrease the FRET efficiency. By adjusting the FRET efficiency of each combination, there is the potential to increase the number of possible probes even further by creating linearly independent combinations.

We conclude that by using an oligo-based approach with internally-labeled fluorophores, we can increase the signal intensity of MuSIC-probe labeled antibodies. MuSIC probe-labeled antibodies may prove useful to increase multiplexing capabilities of full spectrum flow cytometry, and also more broadly where increased multiplexing at single-cell or sub-cellular resolution is needed, including cell-type profiling, tissue studies, and immunofluorescence imaging.

## Acknowledgments

MRB acknowledges funding from Clemson University Creative Inquiry, NIH/NCI Grant R21CA196418, and NIH/NIGMS Grant R35GM141891. MEM received funding from the Department of Education Grant P200A180076. HS acknowledges support from NIH (R01MH081923, 1P20GM130451, R15CA280699) and NSF (CAREER MCB 1749778).

## Methods

### Measuring fluorescent oligo emission spectra

All oligos (Integrated DNA Technologies, Table 1) are resuspended in ddH20 at 100 μM. In a black 96-well plate (Fisher Scientific Cat: 655900), 200 pmols of the fluorescent oligo(s) are added to the well and the volume is brought up to 50μl with PBS. The fluorescent emission spectra are gathered using a Synergy MX microplate reader (Biotek) with parameters set to a slit width of 9 nm, taking readings from the top, an excitation wavelength set to the maximum excitation wavelength for that fluorophore, and an emission wavelength starting 30 nm after the excitation wavelength (Table 2) and emission collected at every nm. The maximum emission intensity was used to quantify results shown in Fig. 2B.

**Table 1.**
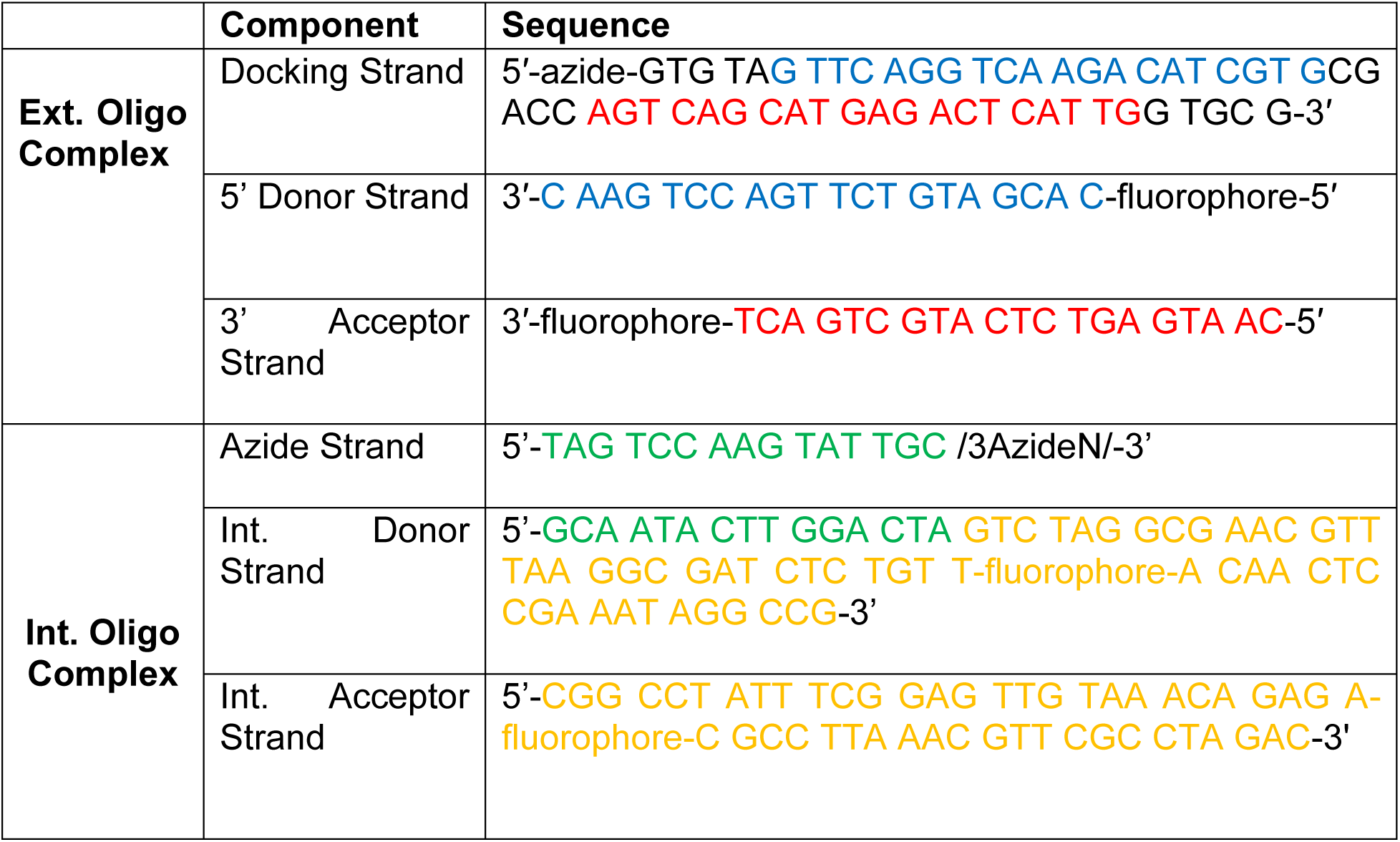
Sequences for the ext. and int. oligo complexes. The text color indicates hybridizing regions.

**Table 2.** Fluorophore modifications for donor and acceptor strands with their corresponding excitation wavelength and emission start wavelength.

| **Fluorophore Modification** | **Excitation (nm)** | **Emission (nm)** |
| --- | --- | --- |
| 6-FAM (Fluorescein) | 490 | 520 |
| Atto 488 | 492 | 522 |
| Atto 532 | 524 | 554 |
| MAX (NHS Ester) | 527 | 557 |
| Cy3 | 534 | 564 |
| Atto 550 | 545 | 575 |
| Tamra (NHS Ester) | 553 | 583 |
| Atto 565 | 561 | 591 |
| ROX (NHS Ester) | 578 | 608 |
| TEX 615 | 583 | 613 |
| Atto 590 | 594 | 624 |
| Atto 633 | 623 | 653 |
| Atto 647 | 632 | 662 |
| Cy5 | 638 | 668 |
| Cy5.5 | 676 | 706 |

**Table 3.** Fluorescence Correlation Spectroscopy, Time-Resolved Fluorescence and Anisotropy Parameters.

|  | Atto488 Ext. Oligo Complex | Atto488 Int. Oligo Complex |
| --- | --- | --- |
| Overall Results |  |  |
| Molecular Brightness (kHz/molecule) | 8.1 | 13.5 |
| Species Average Lifetime ($\langle\tau\rangle_x$) (ns) | 2.58 | 3.05 |
| Diffusion coefficient ($\mu\text{m}^2/\text{s}$) | 100 | 120 |
| Quantum Yield | 50% | 60% |
| Time Resolved Fluorescence and Anisotropy Parameters |  |  |
| $A_1$ | 57% | 68% |
| $t_1$ (ns) | 4.0 | 4.0 |
| $A_2$ | 43% | 32% |
| $t_2$ (ns) | 0.76 | 0.92 |
| $\chi^2$ | 1.65 | 1.64 |
| Rotational Time (ps) | 280 | 350 |
| Fluorescence Correlation Spectroscopy |  |  |
| $G_0$ | 1.00 | 1.00 |
| $N$ | 0.49 | 0.50 |
| $t_{\text{diff}}$ (ms) | 0.71 | 0.59 |
| $\omega_0$ | 4.73 | 4.73 |
| $A_{T_1}$ | 0.28 | 0.26 |
| $t_{T_1}$ ($\mu\text{s}$) | 2.4 | 0.9 |
| $A_{T_2}$ | 0.13 | 0.082 |
| $t_{T_2}$ (ms) | 0.19 | 0.088 |
| $\chi^2$ | 0.96 | 0.86 |

### Labeling Antibodies

Antibodies are conjugated as previously described^13^. In short, the antibody (CD8 clone RPA-T8; Biolegend Cat: 301002; CD4 clone RPA-T4; Biolegend Cat: 300502; CD3 clone UCHT1; Biolegend Cat: 300402) is incubated with DBCO-Peg5-NHS Ester (linker; 10mM in DMSO; Click Chemistry Tools Cat: 1378531-80-6) in 60 molar excess (10 μg of antibody and 2.8 μg of linker) for 30 minutes at room temperature. Post-incubation, the excess linker is removed with Amicon Ultra 100 kDa molecular weight cut-off filters (Fisher Scientific Cat: UFC5100BK). The antibody-linker retentate is collected. Two oligo complexes are created using external (ext.) or internal (int.) fluorophore modifications.

For externally-modified oligos, a 20 bp oligo with a 5’ fluorophore modification (5’ donor strand) and a 20 bp oligo with a 3’ fluorophore modification (3’ acceptor strand) are co-hybridized to a 55 bp oligo with a 5’ azide modification (docking strand) (Integrated DNA Technologies, Table 1) in a 1:1:1 ratio (0.4 nmol of each oligo) to form the ext. oligo complex.

For internally-modified oligos, a 15bp oligo with a 3’ azide modification (azide strand) and a 50 bp oligo with an internal fluorophore modification (int. acceptor strand) are co-hybridized to a 65 bp oligo with an internal fluorophore modification (int. donor strand) (Integrated DNA Technologies, **Table 1**) at a 1:1:1 ratio to one another (0.4 nmol of each oligo) to form the int. oligo complex.

For each, oligo mixtures are incubated for five minutes at room temperature in the dark to allow for complex formation. These complexes (0.4 nmol of each oligo) are then added to the antibody-linker retentate at a 6-molar excess to the original 10 ug of antibody. The volume is brought up to 100 μl with PBS and incubated at 4°C overnight in the dark.

We used int. labeled-antibodies for up to 75 days post-labeling, with storage in PBS at 4°C, with no detectable deterioration in staining properties in downstream flow cytometry assays.

Conventionally-labeled antibodies (anti-CD8 as above) are labeled as per the manufacturer’s instructions (Biotium Mix-n-Stain, Cat: 92446).

### Preparing Peripheral Blood Mononuclear Cells

Normal Peripheral Blood Mononuclear Cells (PBMCs) (Precision for Medicine; 10M cells/vial) are thawed and counted with a hemacytometer. Cells are washed twice with cold (4°C) stain buffer (0.01 g/ml BSA in PBS) at 300 x g for 5 min. Post-wash, the cells are resuspended in cold stain buffer and divided into 100 μl aliquots containing 10^6^ cells. For the experiments in Figures 5 and 6, the same lot of PBMC stocks was used, so similar results could be expected across all replicates.

### Staining PBMCs

In order to block non-specific Fc-mediated interaction,1 μg of normal Rabbit IgG (ThermoFisher Cat: 31235) is added to the cell sample and incubated for 10 minutes at room temperature. Conventionally, ext., and int. labeled-antibodies are made for staining using the protocols described above (10μg of antibody each); (1) CD8 (clone RPA-T8; Biolegend Cat: 301002) labeled with Atto488 ext. MuSIC probes, (2) CD8 (clone RPA-T8; Biolegend Cat: 301002) labeled with Atto488 int. MuSIC probes, (3) CD4 (clone RPA-T4; Biolegend Cat: 300502) labeled with Atto488/647 int. MuSIC probes, (4) CD3 (clone UCHT1; Biolegend Cat: 300402) labeled with Atto647 int. MuSIC probes, and (5) CD8 (clone RPA-T8; Biolegend Cat: 301002) labeled with CF488A (Biotium Cat: 92446). Antibody concentration is adjusted to 0.25 ug/ul for each sample. The labeled antibody is added to the cell sample at the appropriate amount as per manufacturer’s recommendations (2μg CD8 antibody, 0.5μg CD4 antibody, and 0.25μg CD3 antibody/ 10^6^ cells) and allowed to incubate in the dark for 20 minutes on ice. Post-incubation, cells are washed twice with 1 ml of cold staining buffer using 300 x g for 5 min. The final cell pellet is resuspended in 0.5 ml of cold staining buffer.

### Flow Cytometry

Stained PBMC samples are analyzed using a Cytek Aurora spectral flow cytometer. First, unstained PBMCs are assayed with the events to record set to 15,000.

The SpectroFlo software (Cytek) is used to gate single cells (lymphocytes, monocytes, and granulocytes) by forward and side scatter. We then further gate specifically over the lymphocyte population, as typical based on light scattering distributions^38^. Using these same settings, the stained cell samples are assayed. To compare fluorescence intensity between Atto488 and CF488A stained samples we calculate the median intensity of the positively stained cells in the maximum emission channel (B2) using the Spectroflo software. Positively stained cells are defined as cells with a staining intensity above that of the unstained cell samples using a marker gate.

To export raw data, we used FlowJo with “scale” values which were verified to match the original data. The custom unmixing scripts were written in python which are available at https://github.com/birtwistlelab/MuSIC-Antibody-Unmixing and their use described therein. The code is verified to execute in a linux environment (but likely in any). The core function used for unmixing is `nnls` within the `scipy` library. Reference spectra for each dye were generated from the median fluorescence intensity in each channel, calculated from positive singly stained cells. The reference spectra for background autofluorescence were generated from the median fluorescence intensity in each channel, calculated from unstained cells.

### Size Exclusion Chromatography / Multi-Angle Light Scattering (SEC-MALS)

The purpose of SEC was to purify labeled antibody samples prepared as above to provide monomeric antibody conjugates for biophysical characterization below. The approximate retention behavior and molar mass determination of the SEC column and MALS detector (Agilent, AdvanceBio PL1180-3301) was estimated first using Bovine Serum Albumin at 0.5 mg/mL, 30 *μL* injection, and a 0.30 mL/min flowrate using a PBS mobile phase. The MALS instrument (Wyatt Technologies, DAWN 785nm) was normalized, aligned, and broadened using the main peak of the eluent BSA, corresponding to unaggregated BSA (∼5 min retention time). The online concentration was determined using a refractive index detector (Wyatt Technologies, Optilab WREX-08), and we assigned each sample a *dn/dc* of 0.185. We injected the labeled antibody solutions prepared as described above using the same conditions as the BSA experiment. As the approximate absolute molar mass determination via MALS indicated (**Fig. S3**), the chromatogram showed two distinct regions. Eluent corresponding to the first region between 2.5 and 3.5 minutes elution time had an approximate molar mass range of that expected for antibody-oligo conjugates (with a degree of labeling spectrum approximately between 1 and 6). This fraction was collected into vials and preserved for further analysis.

### Fluorescent Correlation Spectroscopy and Time-Resolved Fluorescence and Anisotropy

Freely diffusing samples diluted to sub-nM concentration were analyzed using a custom-built confocal microscope^39^. Samples were excited with a 485 nm pulsed diode laser (LDH-D-C-485, PicoQuant, Germany) operated at 40 MHz. The laser power at the objective was 141 µW. Emission was detected via two PMA detectors (PicoQuant, Germany), allowing for separation into parallel and perpendicular polarization components. A clean-up emission filter (ET525/50, Semrock) is placed before each detector. To ensure temporal data registration of the two synchronized emission channels, we used a HydraHarp 400 TCSPC module (PicoQuant, Germany) in Time-Tagged Time-Resolved mode with a resolution of 1 ps.

Samples were imaged in NUNC chambers (Lab-Tek, Thermo Scientific) that were pre-coated with a solution of 0.01% Tween 20 (Thermo Scientific) in water for 30 min to minimize surface adsorption. Before measurements, chambers were rinsed with buffer to ensure clean measurements. The instrument response function (IRF) was found by measuring water while the protein-free buffer was used for background subtraction. Samples were diluted in charcoal-filtered PBS (10 mM sodium phosphate, pH 7.4, 137 mM NaCl, 2.7 mM KCl) to ∼500pM and measured for 2 minutes.

Software correlations were fit with a 3-dimensional Gaussian with two triplet terms (Eq. 1). The confocal geometric parameter (ω_o_) was determined using Rhodamine 110 as a standard. Diffusion time (t_diff_), molecule count (N), baseline term (*G*_∞_), dark state times (*t*_*T*_1__ and *t*_*T*_2__), and their corresponding fractions (*A*_*T*_1__ and *A*_*T*_2__) were considered as free parameters.

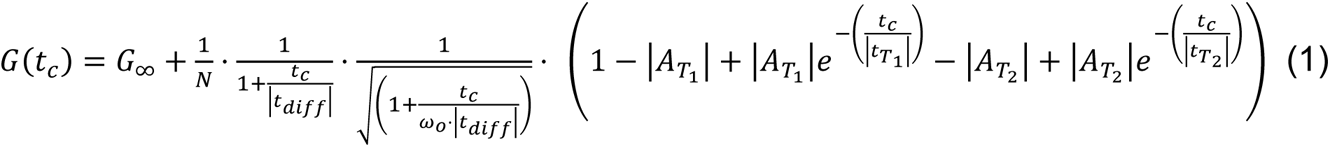

Fluorescence lifetime and rotational correlation times for the samples were found by determining the minimum number of free parameters required to minimize the χ^2^. This was determined to be two fluorescence lifetimes and one rotational time for each sample. The comparison between parallel and perpendicular polarized light about the original laser pulse gives the time-resolved fluorescence anisotropy (r(t) – Eq. 2). The time-resolved anisotropy was fit to Eq. 3.

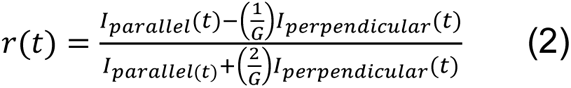

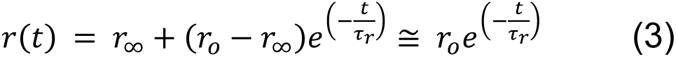

For lifetime fitting, the parallel and perpendicular components were combined using the G-factor (G=1.04) determined using Rhodamine 110 as a standard. Then the fluorescence decays were fit using Eq. 4 with two fluorescence lifetimes for minimizing χ^2^. Then Eq. 5 was used to determine the species average lifetime.

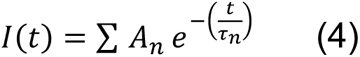

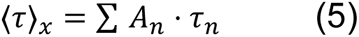

---

For Table of Contents Only

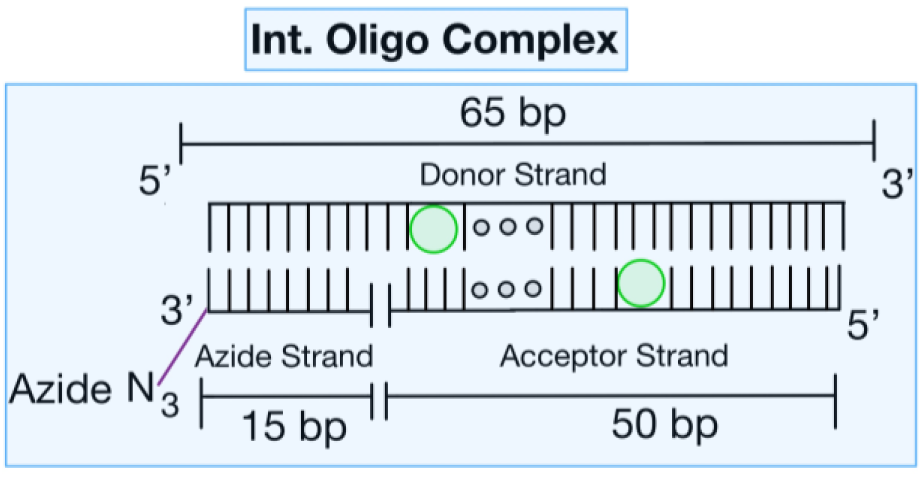

**Figure S1.**
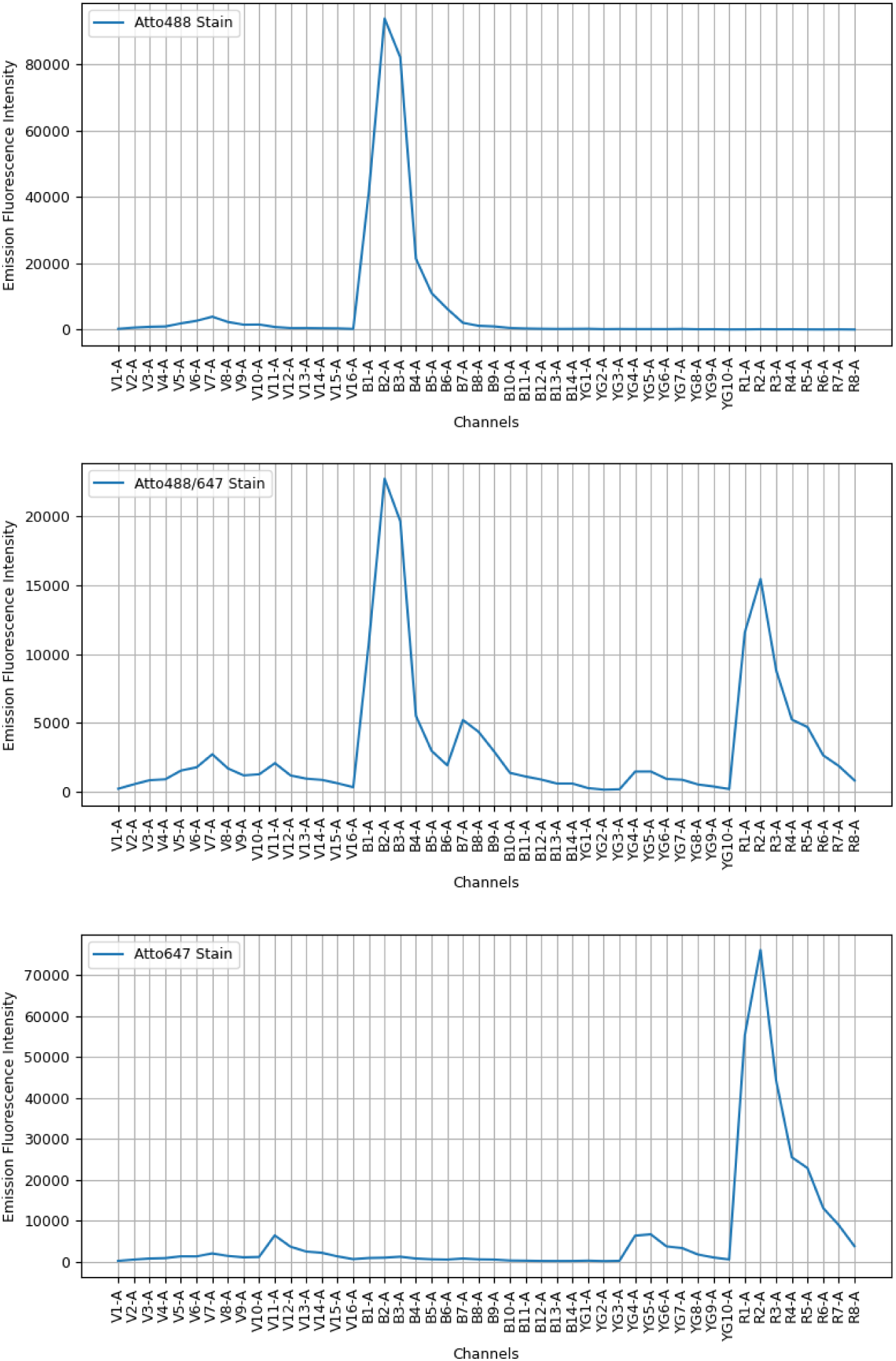
Emission Spectra of the Different Probes. Fluorescence intensities of cells in both unstained and stained-cell groups were measured, and raw data were exported. The median fluorescence intensity (MFI) of each channel in the unstained cells was considered as the autofluorescence. By subtracting this autofluorescence from the corresponding channel’s fluorescence value in the stained cells, the true signal of each stained cell was obtained. The MFI of this true signal of each channel was utilized for spectrum plotting of the stained-cell group.

**Figure S2.**
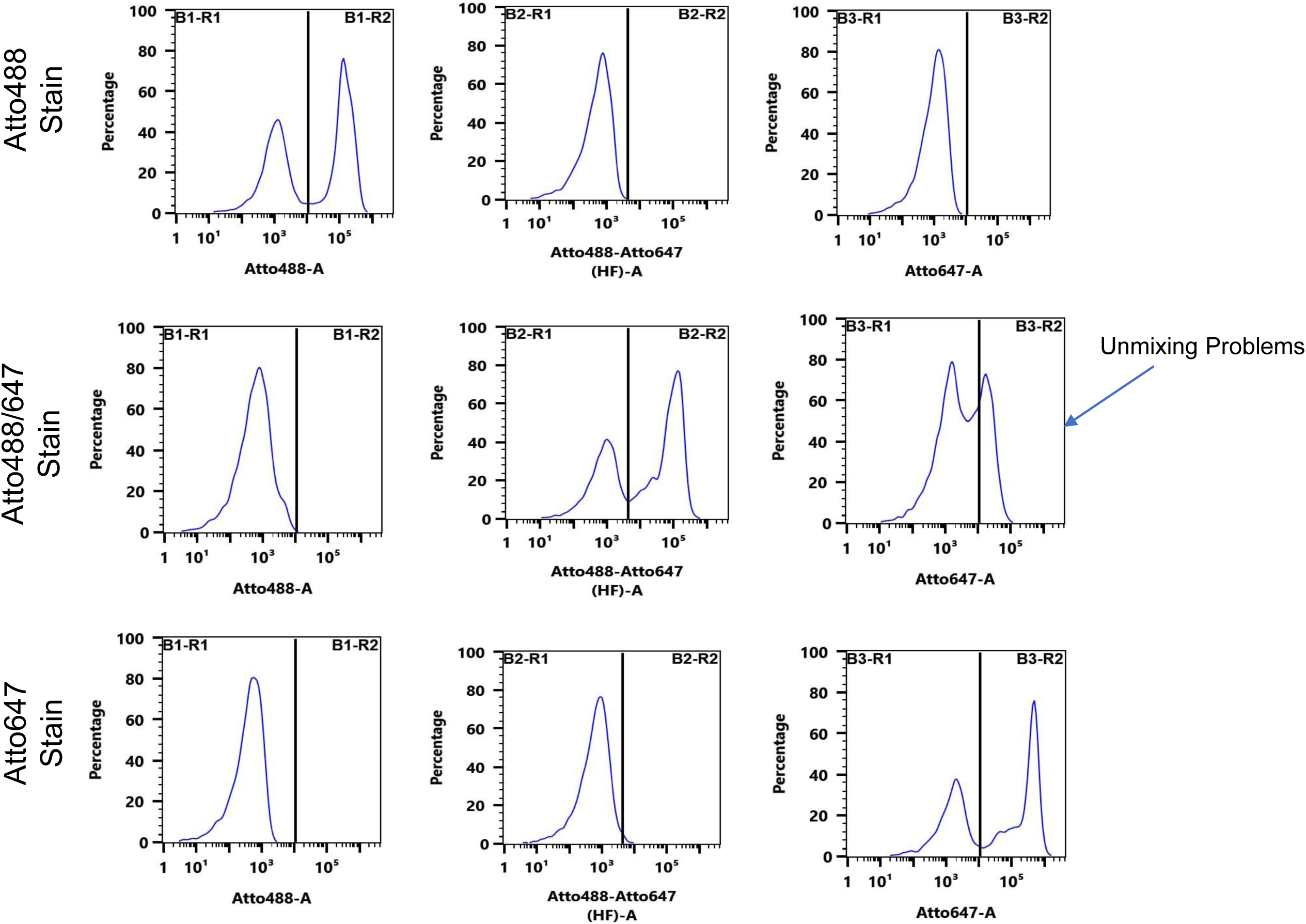
Unmixing Analysis of Single-Stained Cells Using Manufacturer Software. PBMCs were singly-stained as indicated, reference spectra generated, and then subjected to unmixing via the Spectroflo software. The arrow with the legend "unmixing problems" expresses the difficulties unmixing Atto488/647 single stained combination as there is a substantial positive cell population for Atto647, despite the lack of staining with that dye.

**Figure S3.**
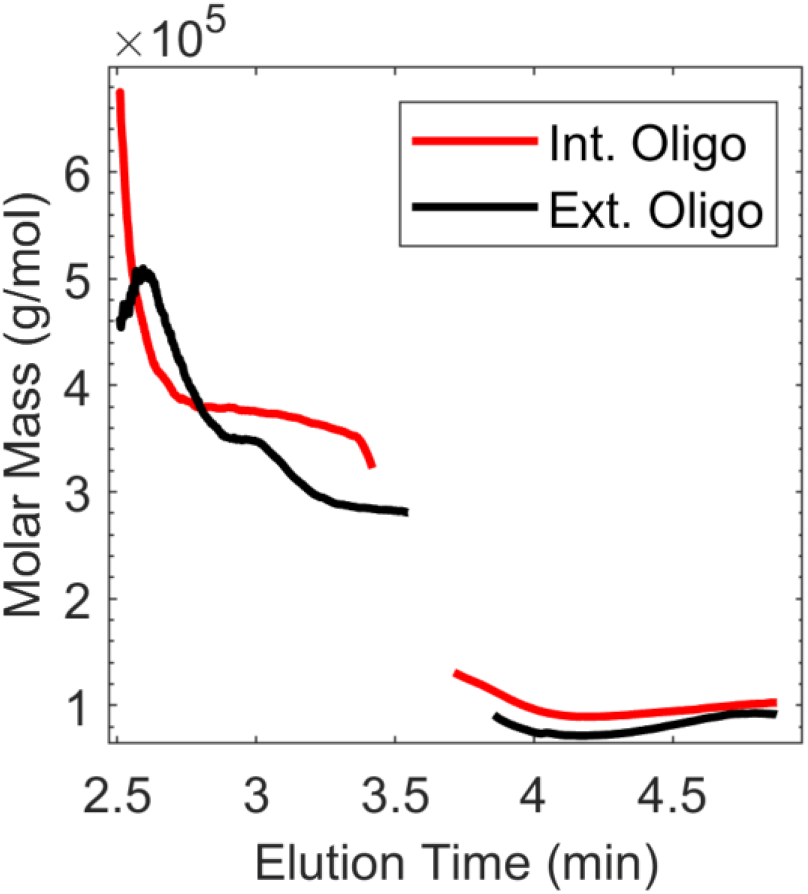
SEC-MALS Data for Labeled Antibody Purification. The molar mass versus elution time measured by SEC-MALS of Int. Oligo labeled antibody (red) and Ext. Oligo labeled antibody (black) solutions. Eluent from 2.5-3.5 minutes was collected for further analysis.

